# Enteric neurons modulate colorectal cancer cell cycle through a PCSK1 - Methionine-Enkephalin Axis

**DOI:** 10.64898/2026.05.03.722515

**Authors:** Philippa Seika, Srinivas Puttapaka, Su Min Hong, Ainslegh Scott, Jared Slosberg, Stefania Bovo Minto, Kevin Haigis, Subhash Kulkarni

**Affiliations:** Department of Surgery, Campus Charité Mitte | Campus Virchow-Klinikum, Charité Universitätsmedizin Berlin, Germany; Division of Gastroenterology, Department of Medicine, Beth Israel Deaconess Medical Center, United States; Dept of Genetic Medicine, Johns Hopkins University, Baltimore, MD, USA; Department of Cancer Biology, Dana-Farber Cancer Institute; Department of Medicine, Harvard Medical School; Harvard Digestive Disease Center; Division of Medical Sciences, Harvard Medical School, United States; Program in Neuroscience, Harvard Medical School, United States

**Keywords:** Enteric neurons, PCSK1, Opioid growth factor receptor, Colorectal cancer, organoids

## Abstract

**Background and Aims:** The tumor microenvironment in colorectal cancer (CRC) is richly innervated, yet the contribution of the enteric nervous system (ENS) to CRC biology remains poorly defined. ENS neurons express proenkephalin (PENK), which can be processed by proprotein convertase 1/3 (PCSK1) to generate Methionine-enkephalin (M-ENK), a bioactive peptide with growth-regulatory potential. We hypothesized that an ENS-derived PCSK1–M-ENK axis restrains CRC proliferation through opioid growth factor receptor (OGFr) signaling and is modulated by stress-associated glucocorticoid receptor (GR) signaling and GLP1 receptor (GLP1R) activity.

**Methods:** Publicly available human CRC single-cell RNA-sequencing datasets were analyzed for OGFr expression. PCSK1 and M-ENK expression in murine ENS and tumor-associated tissue was assessed by immunofluorescence. Functional studies were performed using murine CRC organoids, and primary murine ENS neurons in mono- and co-culture. CRC proliferation was quantified by EdU incorporation following treatment with recombinant M-ENK, recombinant PCSK1, OGFr synthetic ligand naloxone, or PCSK1 inhibitors. Effects of dexamethasone and liraglutide on PCSK1 expression in ENS-containing murine tissue were evaluated.

**Results:** OGFr was enriched in CRC cells and positively associated with *KRAS* gene expression. A subset of adult murine colonic myenteric neurons expressed PCSK1 and M-ENK. M-ENK dose-dependently suppressed proliferation of CRC organoid cells. ENS neurons also suppressed CRC proliferation in a PCSK1-dependent manner. Dexamethasone reduced, whereas liraglutide increased, PCSK1 expression.

**Conclusions:** These findings define a previously unrecognized ENS-derived neuro-oncologic pathway that is associated with reduced CRC cell proliferation and identify the GR/GLP1R–PCSK1–M-ENK axis as a potentially actionable therapeutic node.

**Summary:** This study identifies a neuronal PCSK1 – M-ENK pathway in the ENS that directly suppresses colorectal cancer growth through local OGFr activation, revealing a previously unrecognized neuropeptidergic mechanism of tumor control within the intestinal microenvironment.

## Introduction

Colorectal cancer (CRC) remains a major global health challenge, ranking as the third most commonly diagnosed cancer and the second leading cause of cancer-related mortality worldwide [1, 2]. Despite significant advances in early detection and treatment, the complex and heterogeneous nature of CRC progression continues to limit therapeutic efficacy and patient outcomes [3]. Traditionally viewed as a disease driven primarily by epithelial transformation, mounting evidence underscores the crucial role of the tumor microenvironment (TME) in shaping the hallmarks of CRC [4].

The CRC TME is heterogeneous at the cellular level and given its location in the colonic tissue, it is richly innervated by both extrinsic neurons of the sympathetic and spinal pathways and by the neurons of the enteric nervous system (ENS) [5]. The ENS is the largest collection of heterogeneous neurons and glial cells outside the brain, and thus, provides the bulk of the innervation to various cells of the gastrointestinal (GI) tract to regulate various GI functions [6]. ENS neurons innervate intestinal epithelial cells and regulate their vital functions, including the healthy proliferation of intestinal stem cells [6–8]. This interaction between ENS neurons and mucosal cells extends from health to oncological conditions as well, as prior reports detail the anatomical and molecular crosstalk between ENS neurons and colorectal cancer (CRC) cells [9, 10]. This ENS - CRC crosstalk allows CRC spread by allowing CRC cells to migrate along ENS neuronal fibers, and promotes CRC cell proliferation in a serotonin-dependent manner [9, 10]. However, while existing studies predominantly focus on how the ENS, whether in healthy or dysregulated states (such as through serotonin overexpression), promotes CRC pathology, not much is known about the steady-state mechanisms through which the ENS may restrain CRC proliferation, even after CRC-initiating genetic mutations have arisen. Identifying these molecular pathways is vital for understanding how the healthy ENS neurons naturally suppresses CRC proliferation, how dysfunctions in these pathways may release the ENS-mediated brake on CRC proliferation, and how specific therapeutic strategies can be employed to reinforce this brake to halt cancer progression.

In this report, we identify a novel mechanism through which ENS neurons suppress proliferation of murine CRC cells carrying various driver mutations. We characterized a distinct population of ENS neurons expressing prohormone convertase 1/3 (PC1/3; PCSK1), which catalyzes the conversion of proenkephalin (PENK) to methionine-enkephalin (M-ENK) [11], a ligand for the opioid growth factor receptor (OGFr) expressed by intestinal epithelial cells and overexpressed by CRC cells [12]. We found that the ENS, through this PCSK1 – M-ENK mechanism, suppresses murine CRC cell proliferation. We further found that while an increase in glucocorticoid receptor (GR) signaling, a major stress-induced endocrine signaling pathway [13, 14], suppresses PCSK1 expression, stimulation of glucagon-like peptide 1 receptor (GLP1R) pathway enhanced PCSK1 expression from adult murine colonic myenteric plexus tissue. Thus, the identification of this pathway not only allows us to understand how a healthy ENS suppresses CRC proliferation and how stress-mediated pathways may dysregulate this pathway to increase CRC risk, but it also helps us understand how GLP1R agonists contribute to decreased risk of developing CRC.

Importantly, our data reveal that the anti-proliferative effects of this pathway are not uniform across genetic contexts, but instead exhibit mutation-dependent sensitivity to neuropeptidergic regulation, with *Kras^G12D^*–mutant organoids remaining responsive whereas *Kras^G12V^*–mutant organoids display resistance. These findings identify a previously unrecognized layer of genotype-specific regulation within the tumor microenvironment, in which enteric neuronal signaling exerts differential control over colorectal cancer cell proliferation.

## Materials and Methods

### Analyses of single cell RNA sequencing data of human colorectal cancer (CRC) cells

We mined publicly available single cell RNA sequencing data on 168,672 epithelial (non-malignant and malignant) cells from primary untreated tumors from 34 mismatch repair-deficient (MMRd) and 28 mismatch repair proficient (MMRp) individuals (with an additional lesion collected for 2 individuals) as well as adjacent normal colon tissue for 36 of the individuals generated by the Hacohen group [15] and deposited on the Broad Single Cell data portal (https://singlecell.broadinstitute.org/single_cell/study/SCP1162/human-colon-cancer-atlas-c295?genes=APC&cluster=Epithelial%20cells%20%28tSNE%29&spatialGroups=--&annotation=ClusterFull--group--cluster&subsample=all&tab=distribution). This webportal was used to query the expression of the genes *Ogfr, Kras, Mki67*, and *Lgr5* in various epithelial cell populations of tumor and healthy tissues.

### Analyses of single cell RNA sequencing data of murine colonic enteric neurons

We data-mined publicly available single cell/nucleus RNA sequencing data compilation, generated by combining data from juvenile and adult ENS neurons by the Southard-Smith lab for this study [16]. The data compilation used has been made publicly by the Southard-Smith lab at https://zenodo.org/records/17420912. Only data on colonic myenteric neurons was utilized for assessing the expression of *Pcsk1, Penk* and other genes in various neuronal subpopulations. Graphs were made using UMAP representation of the data using Cellxgene.

### Ex Vivo Whole-Mount Tissue Culture With 500 nM Liraglutide

Prepared colon tissue pieces were incubated under standard tissue culture conditions (37°C, 5% CO₂) in sterile 12-well plates containing stem cell medium (SCM). SCM consisted of neurobasal medium containing containing L-glutamine (Invitrogen), B27 supplement (Invitrogen), and Antibiotic–Antimycotic (Invitrogen), bovine serum albumin (Sigma), and Primocin (Invitrogen). Control tissues were cultured in SCM alone, while experimental tissues were treated with 500 nM liraglutide for 12 hours. Following incubation, tissues were washed in ice-cold PBS. Tissues were snap frozen in liquid nitrogen & stored at −80°C preceding further experimental analysis.

### Cell Culture and Organoid Culture

Murine colorectal cancer organoids, harbouring either *Kras* wild-type (WT) or *Kras* G12D mutations were established by intercrossing conditional mutant alleles of *Apc* (*Apc^lox/+^*) and *Kras* (*Kras^LSL-G12V/+^, Kras^LSL-G12D/+^*) with animals carrying a colonic epithelium-specific Cre transgene (*Fabp1-*Cre). Organoids were maintained in a Matrigel^®^ (Corning, Corning, NY, USA) matrix within a complete organoid medium composed of 47.5 mL DMEM/F12 (Thermo Fisher Scientific), supplemented with 1000 µl B27, 500 µl HEPES, 500 µl Glutamax, 500 µl N2 (X100), 100 µl Primocin, 100 µl N-acetylcysteine, 12.5 µl EGF, and 50 µl Noggin (Thermo Fisher Scientific or R&D Systems).

Organoid cultures were passaged every 3-5 days using the following protocol: Matrigel containing organoids were thawed, and 55 µl of Dispase (Thermo Fisher Scientific) was added per well. After 2 hours of incubation, the content of each well was pipetted up and down and transferred to a 15 ml conical tube. The suspension was centrifuged at 400g for 5 minutes, and the supernatant was discarded without disturbing the pellet. The pellet was then resuspended in 1 ml of TrypLE Express (Thermo Fisher Scientific) and incubated in a bead bath at 37°C for 3 minutes. Following incubation, 1 ml of DMEM with 10% FBS or fresh organoid medium was added, and the suspension was centrifuged again at 400g for 5 minutes. The supernatant was aspirated, and this TrypLE/DMEM+FBS wash step was repeated two more times to ensure complete dissociation. Finally, the dissociated organoids were resuspended in fresh organoid medium, mixed with new Matrigel, and plated for continued culture.

### ENS cell culture

Adult murine enteric neurons were cultured from 2-month-old C57BL/6 mice. Mice were anesthetized with isoflurane and euthanized by cervical dislocation. A laparotomy was performed and the small intestine was removed and lavaged with PBS containing penicillin-streptomycin (PS; Thermo Fisher Scientific), then cut into 2-cm-long segments and placed over a sterile plastic rod. A superficial longitudinal incision was made along the serosal surface and the longitudinal muscle containing myenteric plexus (LM-MP) tissue was peeled off from the underlying tissue using a wet sterile cotton swab and placed in Opti-MEM medium (Thermo Fisher Scientific) containing PenStrep (Thermo Fisher Scientific). For co-culture experiments, LM-MP was dissociated using a digestion buffer comprised of of M199 media (Thermo Fisher Scientific) containing 0.1% BSA, 1 mM CaCl_2_, 20 mM Hepes, 150 μM P188, 50 U/ml DNase I (Worthington), and 1.1 mg/ml collagenase (Sigma) for 40 min at 37 °C and 5% CO_2_. Dissociated cells were then cultured in stem cell media (SCM, comprising of Neurobasal media, B27, BSA and β-mercaptoethanol) containing fibroblast growth factor-β (FGFb), epidermal growth factor (EGF), and glial derived neurotrophic factor (GDNF, all 20 ng/ml, Peprotech) for 5 days to generate neurospheres that contain both differentiated neurons and neural progenitor cells, as performed before [17]. The neurospheres were then co-cultured with murine CRC organoids in SCM without growth factors at 37°C in a humidified atmosphere with 5% CO_2_ for 48 hours.

For experiments involving dexamethasone treatment, neurons were treated with treated with Vehicle (10% DMSO in saline) or dexamethasone (5 µM in 10% DMSO) (Sigma-Aldrich, St. Louis, MO, USA). For experiments involving GLP-1 receptor agonist treatment, ENS neurons were treated with liraglutide (MedChemExpress, catalog HY-P1535), prepared as a stock solution in sterile water and diluted in culture medium to final concentration 500 nM. Cells were exposed to liraglutide for 12 hours under standard culture conditions (37°C, 5% CO₂).

### Cell proliferation assay

Cell proliferation was quantified using the EdU (5-ethynyl-2’-deoxyuridine) incorporation assay, based on the Click-iT™ EdU Cell Proliferation Assay Kit (Thermo Fisher Scientific). For organoid proliferation assays, organoids were dissociated into small clusters and plated at a density of approximately 10,000 cells/clusters per well on coverslips in 24-well plates, embedded in Matrigel. Organoids were allowed to establish for 2-3 days before treatment.

For single-agent treatments, cells/organoids were treated with recombinant Met-enkephalin (M-ENK) (Sigma-Aldrich, St. Louis, MO, USA) naloxone (Sigma-Aldrich, St. Louis, MO, USA) at defined concentrations. For organoid-enteric neuron co-culture experiments, organoids were cultured with primary murine enteric neurons for 48 hours. During this co-culture period, pharmacologic modulation of PCSK1 activity was performed using the PCSK1 inhibitor Decanoyl-RVKR-CMK (Tocris Bioscience, Cat. No. 3501) and recombinant human PCSK1 (OriGene, Cat. No. TP303494). Decanoyl-RVKR-CMK was used at a final concentration of 50 µM, which served as the standard working concentration for all experiments reported in the Results; in selected experiments, a dose range of 25 – 100 µM was additionally tested to assess dose dependency. Recombinant human PCSK1 was used at a final concentration of 10 nM and diluted in assay buffer containing 25 mM MES and 5 mM CaCl₂ (pH 6.0) to maintain enzymatic stability. Enzyme reactions were carried out under conditions ensuring linear substrate cleavage kinetics over the incubation period.

Following treatment, cells/organoids were incubated with EdU at a final concentration of 10 µM for 2 hours at 37°C. Cells/organoids were then fixed with 4% paraformaldehyde (PFA) for 20 minutes and permeabilized with 0.5% Triton X-100 for 30 minutes. EdU detection was performed according to the manufacturer’s instructions using the Click-iT reaction cocktail containing Alexa Fluor 488 azide. Nuclei were counterstained with Hoechst 33342 (Thermo Fisher Scientific) for 30 minutes. For neuronal visualization in co-cultures, cells were additionally stained with PGP9.5 antibody overnight at 4°C, followed by appropriate secondary antibody incubation as described below.

Images were captured using a Leica Stellaris 5 confocal microscope. Proliferation was quantified by counting the percentage of EdU-positive nuclei relative to total Hoechst-stained nuclei in at least 20-30 randomly selected organoids per condition. Image analysis was performed using ImageJ software (NIH, Bethesda, MD, USA) or custom Python scripts primarily leveraging the scikit-image (skimage) library for image processing and segmentation, alongside NumPy and SciPy for numerical operations and data manipulation.

### Tissue preparation

Mice were anesthetized with isoflurane before euthanasia by cervical dislocation. For the isolation of small intestinal tissue, a laparotomy was performed. The entire small intestine was then carefully removed, thoroughly rinsed with PBS containing penicillin-streptomycin (PS; Invitrogen), and subsequently cut into 2-cm-long segments, ensuring that sections from the duodenum, jejunum, and ileum were kept separate. Next, these tissue segments were positioned over a sterile plastic rod. A superficial longitudinal incision was made along the serosal surface, and the longitudinal muscle-myenteric plexus (LM-MP) was gently peeled off from the underlying tissue using a wet sterile cotton swab. The isolated LM-MP was then placed in Opti-MEM medium (Invitrogen) containing Pen–Strep. Following this, the tissue was laid flat and fixed with freshly prepared ice-cold 4% paraformaldehyde (PFA) solution for 4–5 minutes in the dark. After fixation, the tissue was stored in ice-cold sterile PBS with Pen–Strep for subsequent immunofluorescence staining and microscopy.

### Immunochemistry

For murine tissue, the fixed longitudinal muscle-myenteric plexus (LM-MP) tissue underwent a series of washes and incubations. First, the tissue was washed twice in PBS in the dark at room temperature. Subsequently, it was incubated in a blocking-permeabilizing buffer (BPB; 3% BSA with TBST) for 1 hour. This was followed by a 24-hour incubation at 16°C in the dark with shaking at 55 rpm, using the primary antibodies listed in **Table 1**.

**Table 1:**
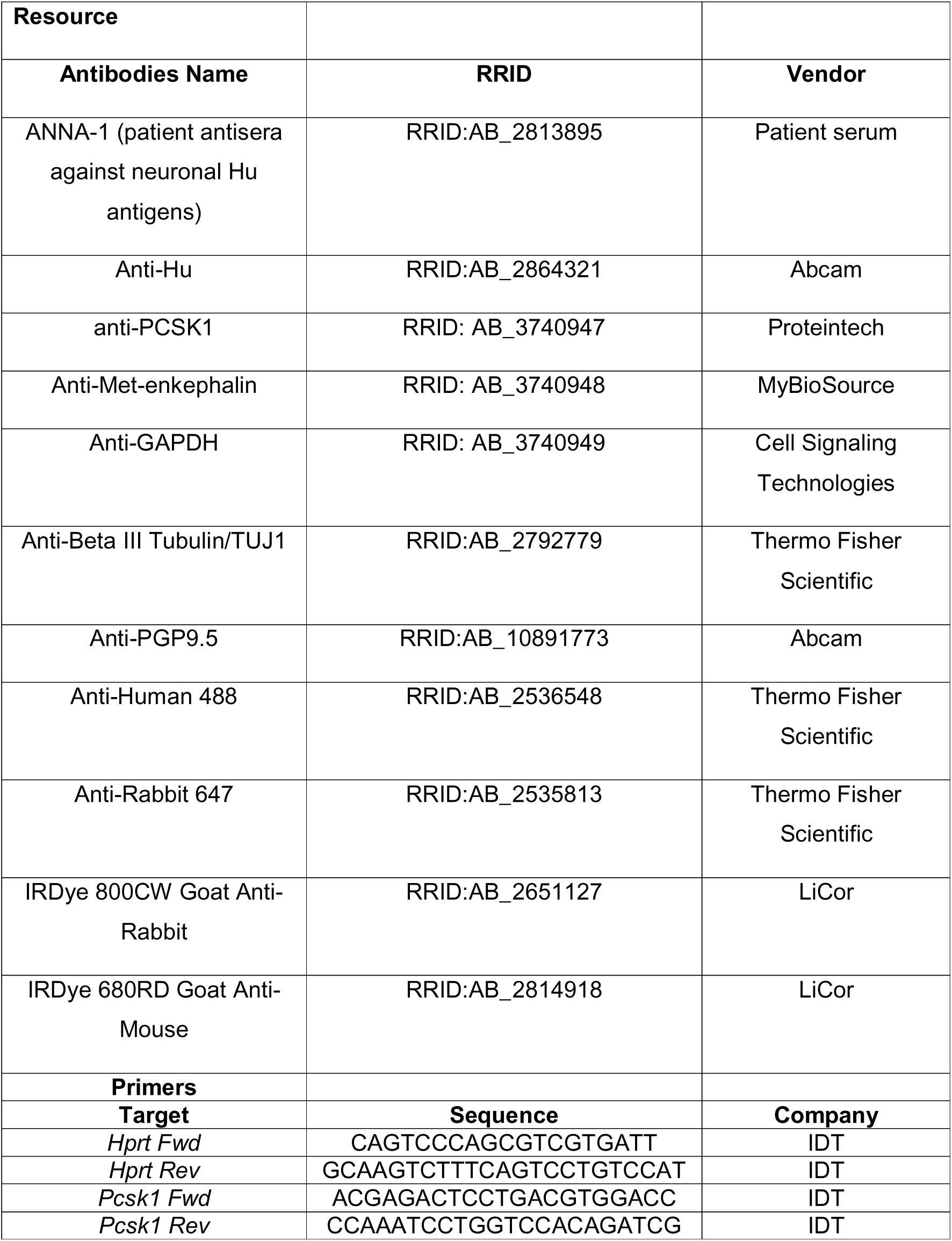

After primary antibody incubation, the tissues were washed three times (15 minutes each wash) in PBS at room temperature in the dark. They were then incubated with the appropriate secondary antibodies (anti-human 488 and anti-rabbit 647; both at 1:500) for 1 hour at room temperature while on a rotary shaker (65 rpm). The tissues were again washed three times in PBS at room temperature, counterstained with DAPI to visualize nuclei, overlaid with ProLong Antifade Gold mounting medium, coverslipped, and finally imaged. The secondary antibody information is listed in **Table 1**.

### Immunofluorescence of FFPE murine tumor sections

Formalin-fixed, paraffin-embedded (FFPE) murine tumor tissues were sectioned at 4 µm thickness. Sections were deparaffinized in xylene and rehydrated through graded ethanol solutions to distilled water. Heat-mediated antigen retrieval was performed using citrate buffer (10 mM sodium citrate, pH 6.0). Sections were allowed to cool to room temperature, washed in PBS, and incubated in blocking–permeabilization buffer (3% BSA in TBST) for 1 hour at room temperature.

Sections were then incubated overnight at 4°C with the following primary antibodies: anti-βIII-tubulin (1:500) and anti-PCSK1 (1:100) (**Table 1**). After washing in PBS, sections were incubated with appropriate fluorophore-conjugated secondary antibodies for 1 hour at room temperature in the dark. Nuclei were counterstained with DAPI. Sections were mounted using ProLong™ Gold Antifade mounting medium, coverslipped, and imaged.

### Protein isolation and western blot analyses

Protein expression of both PCSK1 and Met-enkephalin (M-ENK) was assessed by Western blotting. For protein extraction, previously isolated and flash-frozen LM-MP tissue was used. Tissues were weighed and placed in 1.5 mL locking microcentrifuge tubes containing 6-8 silicone beads. A lysis solution consisting of 1X RIPA buffer supplemented with Halt™ Protease Inhibitor Cocktail (Thermo Fisher Scientific) and Phosphatase Inhibitor Cocktails 2 and 3 (Sigma-Aldrich) was added to each tube. The tissue was homogenized on ice using a bead beater for 5 minutes in a cold room. Following homogenization, the samples were incubated on a shaker for 30 minutes at 4°C to ensure complete lysis. The lysates were then clarified by centrifugation at 12000g for 20 minutes at 4°C. The resulting supernatant, containing the total protein extract, was carefully collected, aliquoted, and stored at −80°C until use. Protein concentrations were determined using the BCA assay (Thermo Fisher Scientific). For M-ENK detection, samples were handled carefully to preserve peptide integrity, including adding the tissues directly to RIPA buffer with protease inhibitors and flash freezing immediately. For western blotting, 30 µg of protein were loaded onto 15% SDS-PAGE gels. For optimal separation and detection of M-ENK, a 16-20% Tris-Tricine gel was used. Proteins were then transferred to PVDF membranes. Membranes were blocked in 5% BSA in TBST for 1 hour at room temperature. Membranes were incubated overnight at 4°C with primary antibodies: anti-PCSK1 antibody (1:1000), anti-M-ENK antibody (1:1000), and anti-GAPDH antibody (1:1000) (**Table 1**). Following incubation, the membranes were washed thrice for 5 minutes each with TBST buffer. They were then incubated with appropriate IRDye-conjugated secondary antibodies (e.g., IRDye 800CW anti-rabbit and IRDye 680RD anti-mouse, LI-COR Biosciences; **Table 1**) in 5% BSA in TBST for 90 minutes at room temperature on a shaker. After three final washes with TBST, the protein bands were visualized and quantified using a LI-COR Odyssey M Imaging System. Densitometric analysis was performed using ImageStudioLite software, and band intensities were normalized to the loading control.

### RNA Extraction, cDNA Synthesis, and Quantitative PCR

Total RNA was isolated from murine LM-MP tissues using the Direct-zol MiniPrep Kit (Zymo Research) according to the manufacturer’s instructions. RNA concentration and purity were assessed spectrophotometrically. 1 microgram of total RNA was reverse-transcribed using SuperScript III Reverse Transcriptase (Invitrogen) in a 20 µl reaction volume. Quantitative real-time PCR (qPCR) was performed using SYBR Green Master Mix on a QuantStudio 3 Real-Time PCR System (Applied Biosystems). Reactions were carried out in 10 µl volumes under the following cycling conditions: initial denaturation at 95°C for 10 min, followed by 40 cycles of 95°C for 10 seconds, 60°C for 10 seconds, and 72°C for 40 seconds. Relative gene expression was calculated using the 2^−ΔΔCt^ method. Expression levels were normalized to the murine housekeeping gene *Hprt*. Primer sequences are detailed in **Table 1**.

### Image Analysis

Image analysis was performed using a custom Python pipeline based primarily on scikit-image. DAPI images were preprocessed with Gaussian smoothing and rolling-ball background subtraction, and nuclei were segmented using Sauvola local thresholding, size filtering, and watershed-based separation of adjacent objects. Organoid regions were identified separately using smoothed image masks with minimum-size filtering. EdU-positive nuclei were quantified from the EdU channel within segmented nuclei. Quantification was performed on all organoids within each image, and values were averaged per image prior to statistical analysis. Code used for analysis is available at https://github.com/klab-ens/imageanalyses_seika.

### Statistical Analysis

All quantitative data are presented as mean ± standard error of the mean (SEM) from at least three independent replicates. Statistical analyses were performed using GraphPad Prism (version 10.0, GraphPad Software, La Jolla, CA, USA). Differences between two groups were analyzed using an two-tailed unpaired Student’s t-test or Welch’s t-test (unequal variance) where appropriate. For comparisons involving three or more groups, one-way analysis of variance (ANOVA) followed by Tukey’s post-hoc test was used. A p-value of less than 0.05 was considered statistically significant. Specific statistical tests used for each experiment are indicated in the figure legends.

## Results

### Expression of Opioid Growth Factor receptor (*OGFR*) is upregulated in human colorectal cancer (CRC) cells

We first tested whether (a) *OGFR* (that encodes Opioid Growth Factor receptor OGFr) is expressed by CRC cells, (b) whether its expression in tumor-derived cells was enhanced in contrast to its expression in healthy cells, and (c) if the expression of *OGFR* was correlated with known marker genes of stemness, proliferation and genes in which mutations were associated with CRC. Our analyses shows that *OGFR* expression was detected in all CRC tumor cells, and that its expression was enhanced in tumor-derived cells relative to control cells (**Fig 1A**). Furthermore, expression of *OGFR* in tumor-derived cells was highest in cells that expressed the stemness gene *LGR5,* and *MKI67*, which encodes for Ki67 – a marker for proliferating cells (**Fig 1A**). *OGFR* expression was also highest in cells that expressed *KRAS*, whose driver mutations are known to be associated with CRC and is a therapeutic target [18] (**Fig 1A**). We next observed that the expression of *OGFR* was better correlated with that of *MKI67* and with *KRAS* in tumor cells than in cells from healthy controls (**Fig 1B, C**) suggesting that *OGFR* is a clinically relevant marker for CRC cells.

**Figure 1:**
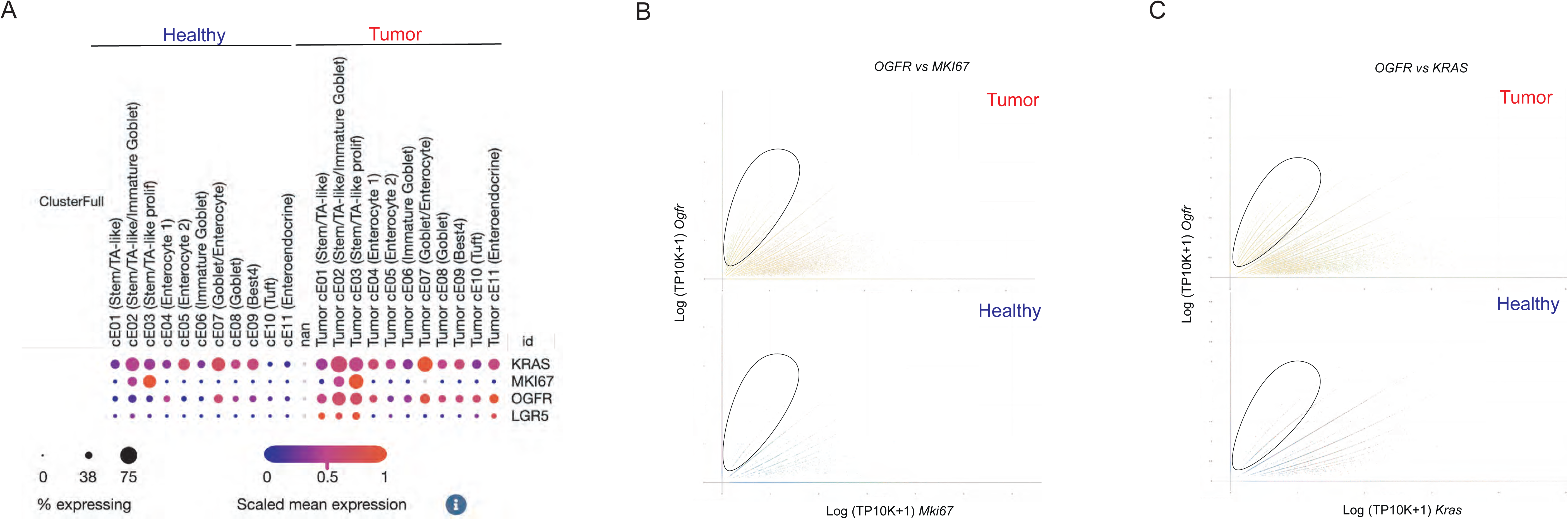
Opioid growth factor receptor (*OGFR*) expression is upregulated in human colorectal cancer cells. (A) Analyses and dot plot-based representation of gene expression of publicly available single cell RNA sequencing data from colonic epithelial cells of healthy patients and of patients with colorectal cancer (CRC) shows that in all CRC cells, and that its expression was enhanced in tumor-derived cells from CRC patients than in control cells. *OGFR* in CRC tissue was highest in *LGR5^+^ MKI67^+^* stem and proliferating cells and in *KRAS^+^* cells. Correlation analyses of (B) *OGFR* versus *MKI67* and (C) *OGFR* versus *KRAS*, stratified by disease state (tumor versus healthy), show that in CRC tumor-derived cells, increasing *MKI67* and *KRAS* expression is associated with higher *OGFR* expression than in cells derived from healthy tissue. The ellipse demarcates the region that shows high *OGFR* expression relative to (B) *MKI67* and (C) *KRAS* in both healthy and CRC tumor-derived cells. The graphs show many more cells within the ellipse in tumor-derived cells than in healthy controls, showing higher *OGFR* expression in *MKI67* and *KRAS*-expressing tumor-derived cells than in healthy cells.

### ENS neurons express Proprotein Convertase Subtilisin/Kexin Type 1 (PCSK1) to generate Methionine-enkephalin (M-ENK)

OGFr is a nuclear receptor that binds to an endogenous pentapeptide Opioid Growth Factor (OGF), which is also known as Methionine-enkephalin (M-ENK) [19, 20]. Pentapeptide enkephalins such as M-ENK (Tyr-Gly-Gly-Phe-Met) and Leucine-enkephalin(L-ENK; Tyr-Gly-Gly-Phe-Leu) are generated by cleavage of proenkephalin (PENK), which is expressed by ENS neurons as well as other GI cells such as fibroblasts [21, 22]. M-ENK, specifically is generated by the cleavage of PENK by the enzyme PCSK1 [23]. We next tested whether ENS neurons express PCSK1 to regulate the generation of M-ENK from PENK. We used a specific anti-PCSK1 antibody and an anti-M-ENK antibody along with anti-Hu antisera (ANNA1) to perform immunostaining of adult murine colonic myenteric neurons and found that that PCSK1 and M-ENK immunoreactivity was observed in subpopulations of colonic neurons (**Fig 2A, B**). We enumerated the abundance of PCSK1-immunoreactive myenteric neurons in the proximal and distal colon and found no significant difference in the abundance of PCSK1-immunoreactive neurons between the two colonic regions (n = 3 mice; total numbers of ganglia/mouse enumerated in proximal colon: 20; mean ± SEM of percentage of PCSK1^+^ neurons/ganglia in proximal colon 16.8 ± 1.6%; total numbers of ganglia enumerated/mouse in distal colon: 15; mean ± SEM of percentage of PCSK1^+^ neurons/ganglia in distal colon 19.9 ± 1.4%; p=0.974; Welch’s t-test (unequal variance)’s t-test **Fig. 2C**). Importantly, we did not detect PCSK1 immunoreactivity in any other cell type in the adult murine colonic longitudinal muscle – myenteric plexus (LM-MP) tissue. In parallel, we have observed that strong PCSK1 immunoreactivity is observed in cells of the human ENS [24]. We further data-mined publicly available single cell/nucleus RNA sequencing data on murine colonic ENS [16] and found that the colonic *Pcsk1*^+^ myenteric neuronal populations comprises of different neuronal subsets, including those that (a) co-express *Vip* and *Nos1*, (b) *Slc17a6* (vGLUT2)^+^ *Calcb* (CGRP)^+^ *Avil* (Advilin)^+^ neurons that may or may not express *Glp1r*, and (c) *Tac1*-expressing neurons (**Suppl. Fig 1**). Interestingly, we observe that *Penk*-expressing neurons are a subset of *Pcsk1*-expressing colonic neurons and a subset of *Penk* and *Pcsk1* co-expressing neurons also express *Glp1r* and the sensory neuronal markers *Avil* and *Calcb*. Since *Calcb* and *Avil* are markers of mucosa-projecting intrinsic primary afferent neurons (IPANs) [25], suggesting that these *Pcsk1^+^ Penk^+^ Glp1r^+^ Calcb^+^ Avil^+^* colonic neurons are IPANs (**Suppl. Fig 1**). These data suggest that colonic enteric neurons are a source of M-ENK, which is produced by cleavage of PENK by neuronally expressed PCSK1, and that PCSK1^+^ M-ENK – releasing neurons can project into the colonic mucosa.

**Figure 2:**
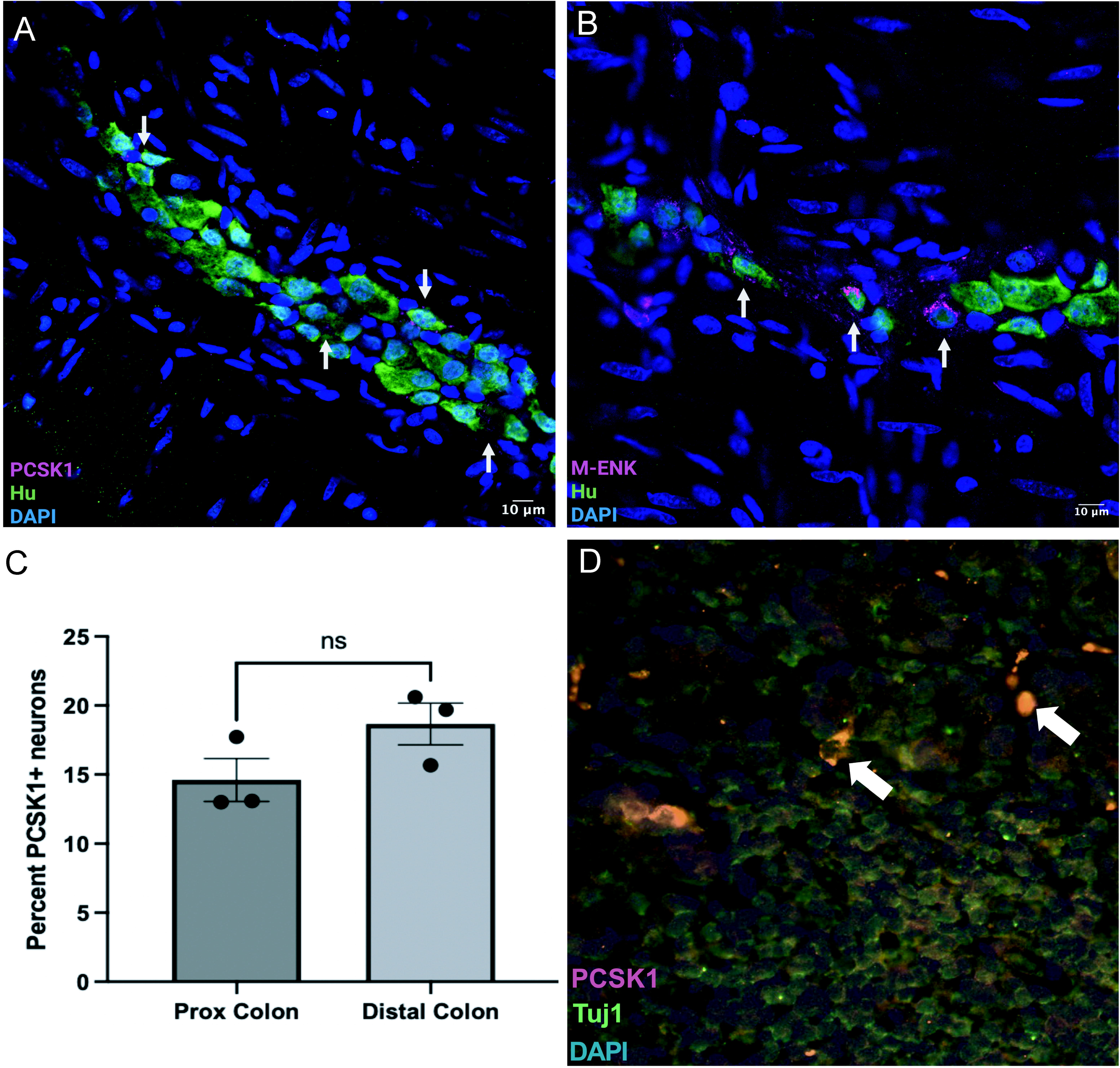
Colonic myenteric neurons express PCSK1 and Methionine-Enkephalin/ Opioid Growth Factor (M-ENK/OGF). Representative immunofluorescence images of adult healthy murine colonic LM-MP tissue immunostained with ANNA1 antisera containing anti-Hu antibodies (green) and with (A) antibodies against PCSK1 (red) and (B) antibodies against M-ENK (red) shows localization (arrows) of PCSK1 or M-ENK with the pan-neuronal marker Hu. Nuclei are stained with DAPI (Blue). Scale bars indicate 10 µm. (C) Quantification of the percentage of PCSK1-immunostained (PCSK1^+^) neurons in murine proximal (n=3) and distal (n=3) colon samples show no significant difference in the proportions of PCSK1-immunostained neurons in the two colonic regions. Data are represented as mean ± standard error of mean; (ns = not significant, Two-tailed unpaired Student’s t-test). (D) Confocal microscopy image showing co-localization (arrows, yellow) of PCSK1 (magenta) and the neuronal marker TUJ1 (green). Nuclei are stained with DAPI (blue).

### PCSK1-expressing neurons project to colorectal cancer (CRC) tumors

To examine neural projections within colorectal tumors, we employed the *Fabp1-Cre:Apc^lox14/+^; Kras^G12D/+^* genetically engineered mouse model, in which Cre-mediated recombination simultaneously induces epithelial *Apc* loss and physiologic activation of endogenous oncogenic *Kras^G12D^* in the colonic epithelium, resulting in invasive adenocarcinoma that recapitulates key features of human CRC [26]. By analyzing formalin-fixed, paraffin-embedded sections using antibodies against the neuronal fiber marker βIII-tubulin (TUJ1) and PCSK1, we found that PCSK1-expressing nerve fibers are present proximate to the CRC tumor site and hence, neuronally-derived PCSK1 and by extension M-ENK constitute a part of the tumor microenvironment (TME) (**Fig. 2D**).

### M-ENK suppresses CRC cell proliferation in a dose- and mutation-dependent manner

Given that M-ENK/OGF is a ligand for OGFr, and OGFr is expressed by proliferating CRC cells, we next tested whether M-ENK affected CRC cell proliferation. For this, we tested whether M-ENK altered cell proliferation in murine CRC organoids, which were generated after introducing defined oncogenic driver alterations in colorectal epithelial cells. The *Kras^WT^* CRC cells retain wild-type *Kras* but harbor alternative oncogenic drivers sufficient to confer a colorectal cancer phenotype, most notably dysregulation of the Wnt/β-catenin pathway, as commonly mediated by loss of APC function. By contrast, cells harboring *Kras^G12D^* and *Kras^G12V^* mutations carry activating point mutations G12D and G12V respectively in the *Kras* gene. We cultured *Kras^WT^*, *Kras^G12D^*, and *Kras^G12V^* murine CRC organoids in complete organoid culture medium containing EdU. Each cell line was treated with either three concentrations of M-ENK (10, 50, or 100 µM), two concentrations of the OGFr ligand naloxone (1 or 10 µM; Sigma-Aldrich, St. Louis, MO, USA), or vehicle alone as control. After culturing for 24 hours in standard tissue culture conditions, we removed the medium and fixed the cells. After performing Click-iT EdU reaction to identify EdU^+^ (cycling) cells, (**Fig 3A**) we calculated the proportions of EdU^+^ cells over all nucleated cells.

**Figure 3:**
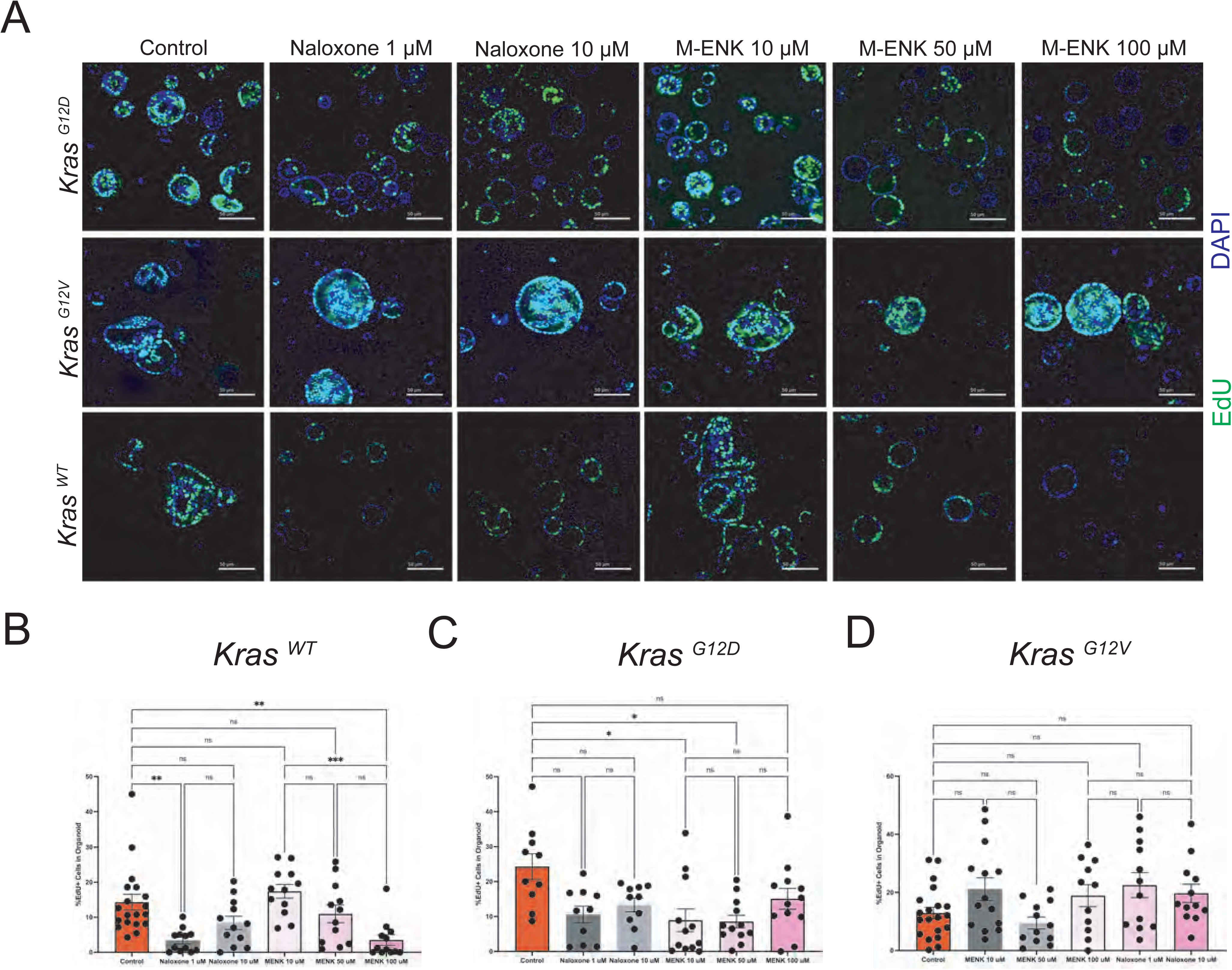
M-ENK suppresses proliferation of murine colorectal cancer cells. (A) Representative confocal images of *Kras^WT^*, *Kras^G12D^*, and *Kras^G12V^*mutations containing murine colorectal cancer cell organoids that were cultured with or without M-ENK or Naloxone in EdU-containing media. After culture, EdU (green) was labelled using Click-iT EdU staining kit to mark proliferating cells and nuclei were stained with DAPI (blue). Scale bars = 100 µm. (B) In *Kras^WT^* mutation carrying cells, Naloxone at lower dose (1 µM) caused a statistically significant reduction in EdU-labeled cells while the Naloxone at higher dose (10 µM) showed no statistically significant effect, when compared to Controls. By contrast, M-ENK at the high dose of 100 µM caused a statistically significant reduction in percentage of EdU-labeled proliferating cells, while it did not cause any statistically significant change in percent of EdU-labeled proliferating cells at the concentrations of 1 µM or 10 µM. (C) In *Kras^G12D^* mutation carrying cells, none of the two Naloxone doses showed any statistically significant effect on altering cell proliferation, when compared to Controls. In these cells, M-ENK at the lower doses of 1 µM and 10 µM caused a statistically significant reduction in percentage of EdU-labeled proliferating cells, while the high dose of 100 µM did not cause any statistically significant change in percent of EdU-labeled proliferating cells. (D) In *Kras^G12V^* mutation carrying cells, neither Naloxone nor M-ENK caused any statistically significant change in percent of EdU-labeled proliferating cells at any concentrations tested. Data are represented as mean ± standard error of mean; (*p < 0.05, **p < 0.01, ***p < 0.001, ns = not significant; one-way ANOVA with Dunnet’s multiple comparisons).

We compared the proportion of EdU⁺ cells per ligand treatment to control using one-way ANOVA followed by Dunnett’s multiple-comparisons test (each concentration vs. control). In *Kras^WT^* organoids, while 10 µM or 50 µM M-ENK treatment did not alter proliferation, 100 µM of M-ENK significantly reduced the proportions of EdU^+^ cells in organoids (n = 19 control, n = 12 per treatment group; mean ± SEM of EdU⁺ cells (%): Control: 14.3 ± 2.2; 10 µM: 17.4 ± 2.0, p = 0.67; 50 µM: 11.0 ± 2.5, p = 0.67; 100 µM: 3.5 ± 1.5, p = 0.0055; one-way ANOVA with Dunnett’s multiple-comparisons; **Fig. 3B**). Similarly, naloxone significantly reduced the proportions of EdU^+^ cells in organoids at the dose of 1 µM. Interestingly, naloxone at 10 µM had no significant effect on the proportions of EdU^+^ cells in organoids (n = 19 control, n = 12 per treatment group; mean ± SEM of EdU⁺ cells (%): Control: 14.3 ± 2.2; 1 µM: 3.4 ± 0.9, p = 0.0037; 10 µM: 8.4 ± 1.9, p = 0.22; one-way ANOVA with Dunnett’s multiple-comparisons; **Fig. 3B**).

In *Kras^G12D^* mutant organoids, the proportions of EdU^+^ cells in organoids at baseline (controls, without any treatment) were (mean ± SEM of EdU⁺ cells) 24.3 ± 3.7%. In these cells, naloxone significantly reduced the proportions of EdU^+^ cells (n = 10 control, n = 10 per treatment group; mean ± SEM of EdU⁺ cells (%): Control: 24.3 ± 3.7; 1 µM: 10.6 ± 2.4, p = 0.020; 10 µM: 13.3 ± 2.0, p = 0.034; one-way ANOVA with Dunnett’s multiple-comparisons test vs. control; **Fig. 3C**). Exposure to M-ENK also significantly reduced the proportions of EdU^+^ cells in organoids at intermediate doses, while the highest dose did not achieve statistical significance (n = 10 control, n = 12 per treatment group; mean ± SEM of EdU⁺ cells (%): Control: 24.3 ± 3.7; 10 µM: 8.9 ± 3.2, p = 0.020; 50 µM: 8.6 ± 1.8, p = 0.003; 100 µM: 15.1 ± 3.0, p = 0.063; one-way ANOVA with Dunnett’s multiple-comparisons; **Fig. 3C**).

Interestingly, by contrast to cells not harbouring *Kras* gene mutations or with *Kras^G12D^* mutations, organoids harbouring *Kras^G12V^*mutations had (mean ± SEM) 12.9 ± 2.0% EdU⁺ cells under control conditions. In these cells, naloxone did not significantly alter the proportions of EdU⁺ cells in organoids (n = 20 control, n = 12 per treatment group; mean ± SEM of EdU⁺ cells (%): Control: 12.9 ± 2.0; 1 µM: 22.5 ± 4.3, p = 0.14; 10 µM: 19.7 ± 3.2, p = 0.20; one-way ANOVA with Dunnett’s multiple-comparisons test vs. control; **Fig. 3D**). Similarly, M-ENK exposure did not significantly affect proliferation at any tested concentration (n = 20 control, n = 14–12–11 per treatment group (10–50–100 µM, respectively); mean ± SEM of EdU⁺ cells (%): Control: 12.9 ± 2.0; 10 µM: 21.2 ± 3.9, p = 0.20; 50 µM: 9.5 ± 2.1, p = 0.26; 100 µM: 18.9 ± 3.8, p = 0.26; one-way ANOVA with Dunnett’s multiple-comparisons test vs. control; **Fig. 3D**).

### ENS neurons modulate CRC organoid cell proliferation in a PCSK1-dependent manner

Given that we found that M-ENK, whose expression is regulated by neuronally-derived PCSK1, significantly reduces proliferation of murine CRC cells, we next investigated whether the presence of ENS neurons exerts a brake on CRC cell proliferation and if so, does the effect manifest in a PCSK1-dependent manner.

For this, we cultured *Kras^G12D^* CRC organoids in 4 conditions in EdU-containing media (complete organoid culture medium): cultured alone without any additional treatments (control group), co-cultured with murine ENS cells, cultured alone in media containing recombinant PCSK1 (10nM), and co-cultured with murine ENS cells in media containing recombinant PCSK1 (10nM). The organoids were cultured for 24h in standard tissue culture conditions, after which the media was removed, the cells were fixed and Click-iT EdU reaction was performed to label EdU^+^ cycling cells (**Fig 4A**). We next enumerated the percentage of EdU^+^ cells over all nucleated cells.

**Figure 4:**
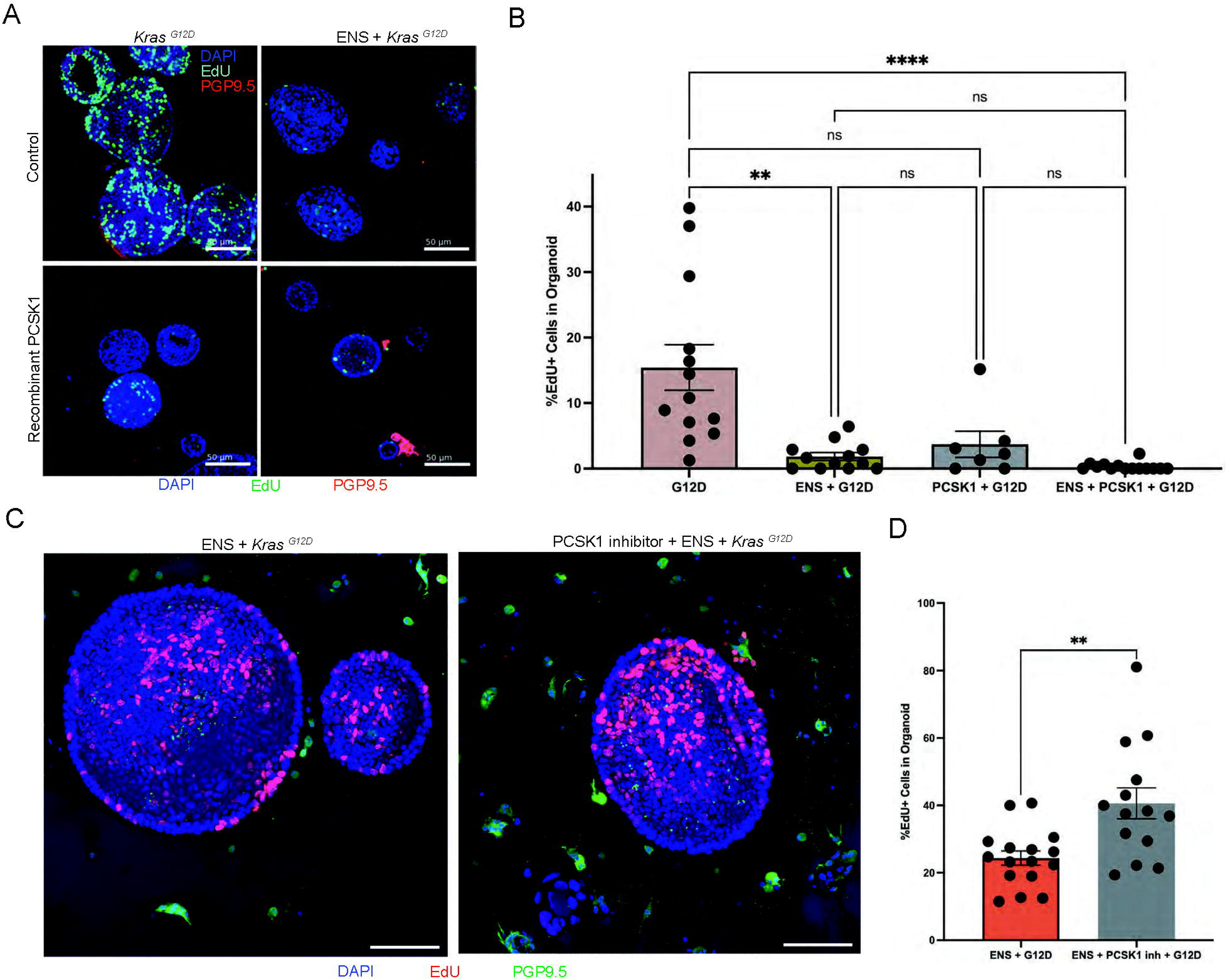
ENS neurons suppress proliferation of colorectal cancer cells in a PCSK1-dependent manner. (A) Representative confocal images of *Kras^G12D^* mutation carrying murine colorectal cancer cell organoids when cultured with or without murine ENS neurons, and with or without recombinant PCSK1 enzyme in EdU-containing media, where after cell fixation, EdU (green) was labelled with Click-iT EdU labelling kit, ENS neurons were immunostained with pan-neuronal antibody PGP9.5 (red) and nuclei were labelled with DAPI (blue). Scale bars = 50 µm. (B) Quantification of percentage of EdU-labeled *Kras^G12D^*mutation carrying cells per total numbers of DAPI-labelled cells in organoids shows that cancer cells when cultured with ENS neurons and when cultured with ENS neurons and PCSK1 show significant reduction in percentage of EdU-labelled proliferating cancer cells in organoids, when compared to controls. Exposure to PCSK1 alone did reduce the percentage of proliferating cells, but this did not meet statistical significance. Data are represented as mean ± standard error of mean; (**p < 0.01, ****p < 0.0001, ns = not significant; one-way ANOVA with Dunnet’s multiple comparisons). (C) Representative confocal images of *Kras^G12D^* mutation carrying murine colorectal cancer cell organoids when cultured with murine ENS neurons, but with or without PCSK1 inhibitor Decanoyl-RVKR-CMK in EdU-containing media, where after cell fixation, EdU (green) was labelled with Click-iT EdU labelling kit, ENS neurons were immunostained with pan-neuronal antibody PGP9.5 (red) and nuclei were labelled with DAPI (blue). Scale bars = 50 µm. (D) Quantification of percentage of EdU-labeled *Kras^G12D^* mutation carrying cells per total numbers of DAPI-labelled cells in organoids shows that cancer cells when cultured with ENS neurons and with PCSK1 inhibitor show significant increase in percentage of EdU-labelled proliferating cancer cells in organoids, when compared to those cultured with ENS cells but without PCSK1 inhibitor. Data are represented as mean ± standard error of mean; (**p < 0.01; Two-tailed unpaired Student’s t-test).

By performing one-way ANOVA with Dunnett’s multiple comparisons test vs. control multiple-comparisons test, we found that, compared with control organoids, CRC organoid proliferation was significantly reduced in the presence of ENS cells (Control vs. ENS: p = 0.0042) and most strongly under combined ENS + PCSK1 exposure (Control vs. ENS + PCSK1: p < 0.0001), while recombinant PCSK1 alone did not significantly reduce proliferation (Control vs. PCSK1: p = 0.1166). ENS co-culture and PCSK1 exposure each reduced proliferation, with the strongest suppression observed under combined conditions (n = 13 for Control group, n = 12 for ENS group, n = 7 for PCSK1 group, n = 14 for ENS + PCSK1 group; mean ± SEM of EdU⁺ cells (%): Control: 15.4 ± 3.5; ENS: 1.84 ± 0.60; PCSK1: 3.72 ± 2.00; ENS + PCSK1: 0.32 ± 0.17; one-way ANOVA with Dunnett’s multiple-comparisons test vs. control; **Fig. 4B**).

These findings demonstrate that recombinant PCSK1 contributes to the anti-proliferative effect of ENS co-culture, suggesting that ENS-mediated suppression of CRC proliferation occurs, at least in part, via PCSK1-dependent mechanisms.

To directly test this dependency, we performed a separate experiment in which ENS co-cultured *Kras^G12D^* CRC organoids were treated with the PCSK1 inhibitor Decanoyl-RVKR-CMK (50 µM) for 24 hours (**Fig. 4C)**. Statistical significance was assessed using an unpaired two-tailed unpaired Student’s t-test. PCSK1 inhibition significantly increased proportions of EdU⁺ cells when compared to Control, demonstrating that ENS-mediated growth suppression requires PCSK1 activity. These data show that neuronally-derived PCSK1 is an important regulator of CRC cell proliferation (n = 15 for Control group, n = 13 for PCSK1 inhibitor group; mean ± SEM of EdU⁺ cells (%): Control: 33.9 ± 2.8; PCSK1 inhibitor: 45.2 ± 3.9, p = 0.009; Two-tailed unpaired Student’s t-test; **Fig. 4D**).

### Glucocorticoid receptor signaling regulates ENS PCSK1 expression and reduces M-ENK

Increased glucocorticoid signaling driven by heightened cortisol release is a hallmark of stress-induced endocrine signaling in the periphery that has been shown to impact the ENS [13, 14]. Stress is known to increase the risk of developing cancers, including CRC [27]. Stress-induced increase in cortisol engages the glucocorticoid receptor, and stress induced increased glucocorticoid receptor (GR) signaling facilitates the effect of stress at a cellular level [28]. Using our *in vitro* system supplemented with a GR agonist dexamethasone to mimic the effect of stress-induced glucocorticoid signaling on ENS – CRC organoids, we tested whether dexamethasone-induced increased GR signaling alters the neuronal PCSK1 – M-ENK pathway. After culturing adult murine colonic longitudinal muscle–myenteric plexus (LM–MP) tissue with or without dexamethasone (5 µM) for 24 hours in Neurobasal medium containing B27 and BSA [14], total protein was isolated and the relative abundance of PCSK1 was assessed by Western blot analysis, normalized to the housekeeping protein GAPDH. Comparison of PCSK1 protein levels between treatment groups revealed a region-specific effect of dexamethasone exposure. While PCSK1 abundance in proximal colon LM–MP tissue was not significantly altered (n = 3/group; mean ± SEM of arbitrary units: Control: 14,086 ± 729; Dexamethasone: 11,402 ± 2,815; p = 0.44, Two-tailed unpaired Student’s t-test; **Fig. 5A**), dexamethasone treatment significantly reduced PCSK1 levels in distal colon LM–MP tissue (n = 3/group; mean ± SEM of arbitrary units: Control: 22,323 ± 1,648; Dexamethasone: 15,104 ± 597; p = 0.036, Two-tailed unpaired Student’s t-test; **Fig. 5B**).

**Figure 5:**
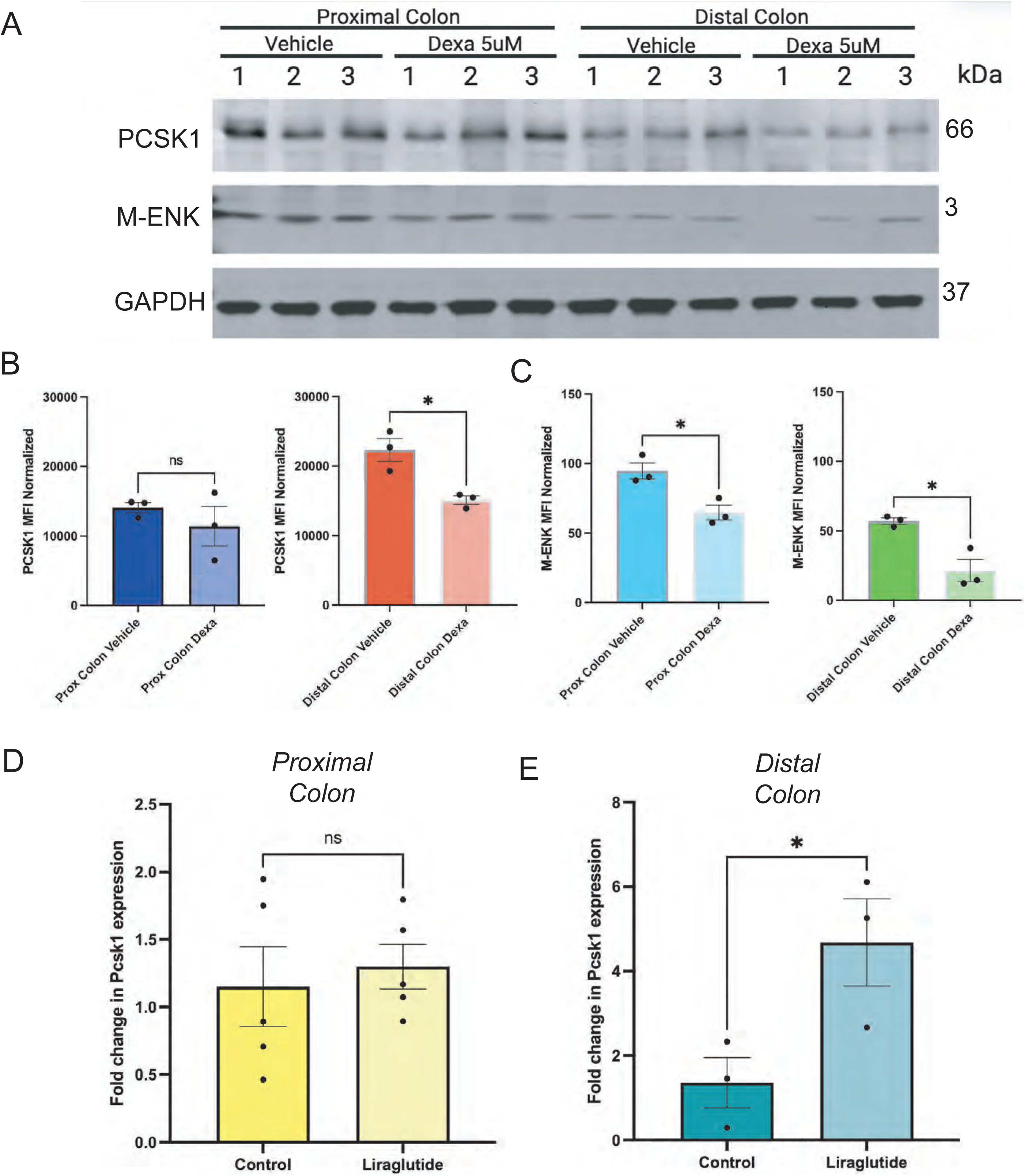
Glucocorticoid receptor signalling and GLP1 receptor signaling have opposite effects on PCSK1 – M-ENK pathway in the colonic ENS. (A) Western Blot images showing protein expression of PCSK1 (66 kDa), Met-enkephalin (M-ENK, 3 kDa), and GAPDH (37 kDa) as a loading and normalization control, in total proteins isolated from adult murine proximal and distal colon LM-MP tissues (from n = 3 mice) that were exposed to Dexamethasone (5 µM) or Vehicle. (B) Graphical representation quantifying normalized PCSK1 protein levels by calculating mean fluorescent intensity (MFI) of PCSK1 normalized to MFI of GAPDH from the Western Blot data presented in (A) shows that Dexamethasone exposure causes a significant reduction in PCSK1 abundance in Distal but not in Proximal colon tissue. Data are presented as mean ± standard error of mean (*p < 0.05, ns = not significant; Two-tailed unpaired Student’s t-test). (C) Graphical representation quantifying normalized M-ENK protein levels by calculating mean fluorescent intensity (MFI) of M-ENK normalized to MFI of GAPDH from the Western Blot data presented in (A) shows that Dexamethasone exposure causes a significant reduction in M-ENK abundance in both Proximal and Distal colon tissue. Data are presented as mean ± standard error of mean (*p < 0.05, ns = not significant; Two-tailed unpaired Student’s t-test). Quantiative RT-PCR (qRT-PCR)-based analyses of *Pcsk1* gene expression, normalized to *Hprt* expression, in adult murine (E) proximal and (F) distal colonic LM-MP tissue cultured *ex vivo* with and without Liraglutide for 16 hours shows that Liraglutide exposure causes a significant increase in *Pcsk1* expression in (F) distal colonic LM-MP tissue but not in (E) proximal colon LM-MP tissue. Data represented as mean ± standard error of mean; (*p < 0.05, ns = not significant; Two-tailed unpaired Student’s t-test).

We next assessed whether dexamethasone-induced suppression of PCSK1 was accompanied by a corresponding reduction in methionine-enkephalin (M-ENK) levels. Western blot quantification of M-ENK, normalized to GAPDH, demonstrated that dexamethasone treatment significantly reduced M-ENK abundance in both regions (Proximal colon LM–MP: (n = 3/group; mean ± SEM of arbitrary units: Control: 94.7 ± 5.8; Dexamethasone: 64.7 ± 5.3, p = 0.019; Two-tailed unpaired Student’s t-test; Fig. 4A, 4C). Distal colon LM–MP: (n = 3/group; mean ± SEM of arbitrary units: Control: 57.1 ± 2.2; Dexamethasone: 21.5 ± 8.1, p = 0.040; Two-tailed unpaired Student’s t-test; **Fig. 5C**).

Together, these data establish that GR signaling negatively regulates the neuronal PCSK1–M-ENK axis, suggesting a potential mechanistic link between psychological and systemic stress signaling and impaired PCSK1–M-ENK biology in the neuro-oncological axis relevant to CRC risk.

### GLP-1 receptor signaling regulates neuronal PCSK1 expression

Recent studies have identified a protective role of glucagon-like peptide-1 receptor (GLP-1R) agonist drugs in CRC [29–31]. Given that GLP-1 receptor is expressed by colonic ENS neurons [32], we hypothesized that this protective effect of GLP-1R agonists against CRC is mediated, at least in part, by GLP-1R–dependent upregulation of PCSK1 – M-ENK biology in colonic ENS neurons. To test this, adult murine proximal and distal colonic LM-MP tissues were again cultured in completely defined medium without (control) and with the GLP-1R agonist liraglutide (500 nM) for 12 hours. After harvesting the tissues, the total RNA and generated cDNA was used to perform quantitative RT-PCR with primers against *Hprt* (housekeeping gene) and *Pcsk1*. We found that while *Pcsk1* expression in proximal colon LM–MP tissue was not significantly altered by liraglutide exposure (n=3/group; mean ± SEM fold change in relative abundance of *Pcsk1* transcript: Control: 1.13 ± 0.32; Liraglutide: 1.30 ± 0.18; p = 0.67, Two-tailed unpaired Student’s t-test; **Fig. 5D**), liraglutide significantly increased *Pcsk1* expression in distal colon LM–MP tissue (n=3/group; mean ± SEM fold change in relative abundance of *Pcsk1* transcript: Control: 1.36 ± 0.41; Liraglutide: 4.68 ± 1.19; p = 0.0497, Two-tailed unpaired Student’s t-test; **Fig. 5E**). Given that we have shown that modulation of PCSK1 abundance regulates M-ENK abundance, we infer that GLP-1R agonism upregulates the neuronal PCSK1–M-ENK axis, which our earlier data shows would then act on OGFr in tumor cells of CRC to suppress their proliferation.

## Discussion

Our study identifies a previously unrecognized neuro-oncogenic mechanism mediated by PCSK1 - Met-enkephalin (M-ENK/OGF) – OGFr axis, through which ENS neurons exert a brake on CRC cell proliferation and hence tumor growth. While earlier studies proposed a pro-oncogenic role of ENS neurons on CRC [9, 10], ours is the first report that highlights a tumor-suppressive neuro-oncological mechanism between ENS neurons and CRC. Our study establishes Opioid Growth Factor receptor (OGFr) as a clinically relevant marker for CRC cells and it shows that neuronally-derived protein convertase 1/3 (PCSK1) enzyme is a key biomolecule that regulates the abundance of OGF/M-ENK that acts on OGFr to reduce CRC cell proliferation. Furthermore, we show that this tumor-suppressive neuronal mechanism is responsive to stress, where stress-driven glucocorticoid signaling suppresses and GLP-1R signaling enhances this tumor suppressive mechanism in the distal colon. These data together suggests how the healthy ENS suppresses CRC proliferation and how physiological and psychological stress may promote CRC cell proliferation in an ENS-dependent manner. We also observe that GLP-1R signaling promotes the expression of the PCSK1 from ENS neurons of the distal colon, suggesting that it would increase M-ENK/OGF levels to suppress CRC cell proliferation in an OGFr-dependent manner. Since GLP-1 release from colonic enteroendocrine cells is known to occur in response to microbe-derived short chain fatty acids (SCFA) [33], these data provide a possible mechanism through which healthy gut microbiota suppress tumorigenesis in the colon [34].

In this study, by mining publicly available single cell transcriptomic data on human CRC cells, we first showed that OGFr is a clinically relevant marker in CRC cells, as its expression is enhanced in tumor cells relative to control healthy cells, and its expression is positively correlated with expression of cell cycle marker *Mki67* (coding for Ki67) and the expression of CRC relevant oncogene *Kras*. Furthermore, given that OGFr signaling has been shown to restrict cell proliferation in other cancers [35], its relevance to CRC was important to test. Since OGF/M-ENK, whose biology is significant to the ENS [36], we hypothesized and tested that M-ENK was indeed generated in the ENS, and that *Penk* and *Pcsk1* co-expressing (and thus M-ENK releasing) myenteric neurons projected into the colonic mucosa. By identifying a subset of neurons that expressed the PC1/3 enzyme PCSK1 that regulates conversion of proenkephalin (PENK, which is expressed by both ENS neurons and intestinal fibroblasts) [22] and that these neurons project to the tumor microenvironment, we show that ENS neurons play a crucial role in regulating CRC biology through PCSK1 – OGF – OGFr signaling.

Using *in vitro* culture of murine CRC cell organoids harboring clinically relevant CRC driver mutations in *Apc* and *Kras* genes, our study establishes how increasing OGFr signaling through known OGFr ligands M-ENK and naloxone suppress CRC cell proliferation. We further established a CRC – ENS co-culture platform through which we interrogated the effect of ENS cells on CRC cell proliferation. We observed that presence of ENS cells suppresses CRC cell proliferation, an effect that is enhanced by supplementing this co-culture system with PCSK1 protein. We further observed that addition of a PCSK1 inhibitor weakened this brake on CRC cell proliferation. Thus, these data establish that neuronally-derived PCSK1 is a key regulator molecule in the suppression of CRC proliferation.

Importantly, we observed that while OGFr signaling exerted a brake on cell proliferation in the absence of *Kras* mutations and in the presence of *Kras^G12D^* mutations, CRC cells harboring *Kras^G12V^* mutations are resistant to the OGFr-mediated brake on cell proliferation. In light of clinical data showing that CRC patients with *Kras^G12V^* mutations have significantly worse overall survival [37] and preclinical data from our group showing that molecular phenotypes of *Kras^G12D^ and Kras^G12V^* mutations are unique [38], the resistance of CRC cells with *Kras^G12V^* mutations to OGFr signaling may help explain the poor prognosis of patients with *Kras^G12V^*mutations.

The expanded knowledge of neuronal regulation of CRC biology also helps us study how factors that are known to increase or decrease CRC risk may exert their effects. In this context, our data on increased glucocorticoid receptor (GR) signaling in ENS, which we have shown occurs as a result of increased psychological stress [14], on reduced PCSK1 and M-ENK abundance may help explain mechanistically how stress increases risk of colorectal tumorigenesis [39]. Similarly, our data on liraglutide’s effect to increase expression of *Pcsk1* may help explain the mechanism through which the observed protective effect of GLP-1R agonist against CRC tumorigenesis [29, 30]. Intriguingly, the effect of dexamethasone and of liraglutide on modulation of PCSK1 occurs only in distal colon and not in proximal colon, suggesting significant differences in the PCSK1-expressing neuronal populations in the two colonic regions. However, we currently have no clarity on the differences in molecular taxonomy of the PCSK1-expressing myenteric neurons in proximal vs distal colon, this will form an important line of enquiry to better understand the translational significance of this pathway.

Another key observation is the partial overlap of PCSK1 and M-ENK expression within a subset of myenteric neurons in the murine colon. This suggests a functional association, but the lack of complete co-localization precludes definitive assignment of M-ENK secretion to PCSK1-expressing neurons. Furthermore, since M-ENK precursor proenkephalin (PENK) is also expressed by fibroblasts, it suggests that neuronally-restricted PCSK1 is the rate limiting step in the genesis of M-ENK and in the neuronal brake on CRC cell proliferation.

An important question remains that if OGFr signaling restricts CRC cell proliferation, then why OGFr expression is enhanced on CRC cells. One possible explanation is that increased expression of OGFr in CRC cells shifts the stoichiometry of M-ENK/OGF – OGFr signaling in favor of OGFr and thus steady state amounts of M-ENK/OGF is insufficient for continuing to exert the brake on CRC cell proliferation. In this scenario, therapeutic interventions that increase M-ENK/OGF levels in the colonic tissue would serve as the backbone of novel interventions to target CRC.

Despite these insights, several limitations warrant consideration. First, the majority of functional experiments were performed *in vitro* using murine organoid and co-culture systems, which cannot fully recapitulate the complexity of the *in vivo* tumor microenvironment, including immune, stromal, and vascular interactions. Accordingly, the extent to which the neuronal PCSK1–M-ENK axis modulates tumor growth *in vivo* remains to be established.

Second, while our data support a role for M-ENK–mediated signaling in regulating CRC proliferation, the precise receptor mechanisms remain incompletely defined. Although we focus on OGFr as a candidate mediator, we cannot exclude contributions from classical opioid receptors or alternative signaling pathways, particularly given the pharmacologic effects observed with naloxone.

Third, the upstream mechanisms governing PCSK1 release from enteric neurons and the downstream intracellular pathways activated by OGFr signaling in CRC cells remain to be elucidated. Addressing these questions will be essential to fully define the therapeutic potential of this neuropeptidergic axis. Our future work will seek to validate these findings in in vivo murine models and with human cells, to determine the net effect of this pathway on tumor growth and metastasis. Furthermore, how GLP-1 generation and release are altered in colonic L-cells and how these mechanisms relate to increased oncogenic potential in colorectal tissues are yet to be elucidated.

In summary, we describe a novel neuro-oncogenic regulatory circuit in CRC, in which enteric neurons, via PCSK1, modulate tumor proliferation through M-ENK and OGFr signaling. These findings provide a mechanistic framework for understanding context-dependent effects of PCSK1, reveal a potential tumor-suppressive function of the ENS, and suggest that therapeutic modulation of the neuronal PCSK1-M-ENK-OGFr axis may represent a previously unappreciated strategy for CRC treatment.

## Acknowledgements

This work was supported by funding from NIA R01AG066768, R21AG072107, and Pilot grant from the Harvard Digestive Disease Core (SK). This work was also supported by funding from the Walter Benjamin Fellowship (528835020) from Deutsche Forschungsgemeinschaft (PS).

## Supplemental Data

**Supplementary Figure 1: Molecular taxonomy of PCSK1-expressing colonic murine myenteric neurons.** Publicly available single cell/nucleus RNA sequencing data on murine colonic ENS [16] was obtained and mined to characterize *Pcsk1*-expressing colonic myenteric neurons. Using data specifically on colonic myenteric neurons, we established that in the UMAP plots (A) *Nos1* and (B) *Chat* are expressed on distinct supersets of neuronal cells. (C) We establish that two separate clusters within the *Nos1+* neuronal set express *Vip,* which we defined with a region of interest (ROI) in green. (D) We established a cluster of *Tac1^+^* neurons within the *Chat^+^* neuronal set and labeled it with a ROI in red. Next we defined (E) 3 clusters with blue ROI that express sensory marker *Avil*, of which one of the clusters (F) express *Slc17a6* (vGlut2), labeled in grey ROI. (G) Two clusters of neurons expressing *Scgn* express *Vip*, as defined by the *Vip*-specific green ROI. (H) *Penk* expression occurs in two distinct clusters, one defined by *Slc17a6*-specific grey ROI and another by *Tac1*-specific red ROI. (I) *Calcb*-expression is found in the *Avil*-expressing clusters, defined by the blue ROI. (J) *Glp1r*-expression is found in two distinct clusters, defined by the orange ROI. Taking together, *Pcsk1-*expression is found in diverse and distinct neuronal subsets, including those that (a) defined by red RO1 and hence are *Tac1*^+^; defined by green ROI and hence express *Vip* and *Nos1*; defined by orange ROI and hence express *Glp1*r, and defined by grey and blue ROIs and hence express *Slc17a6* and *Avil/Calcb*. (L) Finally, we observe that expression pattern of *Pcsk1* and its subset-defining ROIs are similar to the expression of choline transporter gene *Slc5a7* in the murine colonic myenteric plexus neurons.

