## Supplementary figures and images for "Enteric neurons modulate colorectal cancer cell cycle through a PCSK1 - Methionine-Enkephalin Axis"

### Suppl. Fig 1

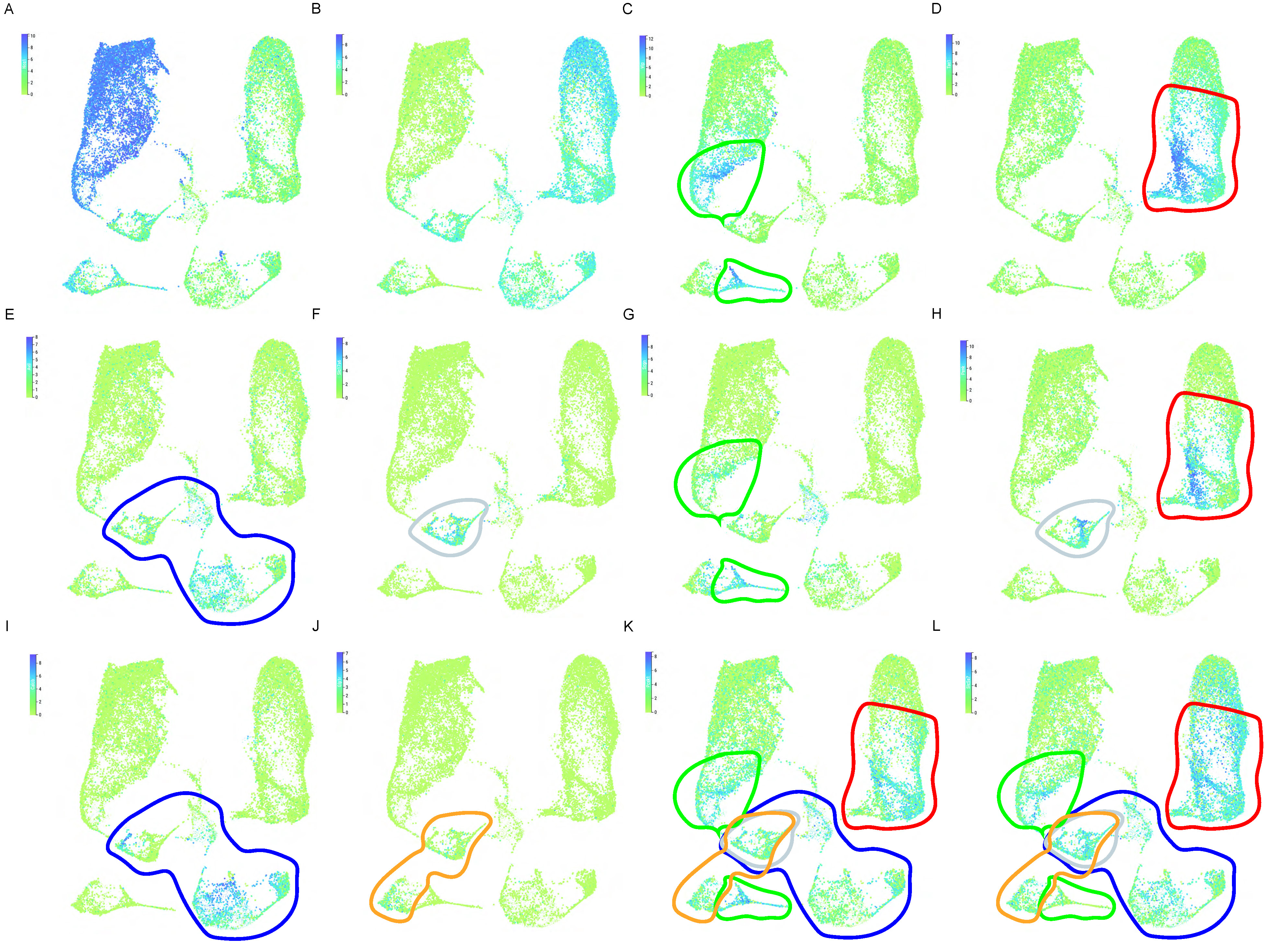
